# Denoising complex-valued diffusion MR images using a two-step non-local principal component analysis approach

**DOI:** 10.1101/2024.10.30.621081

**Authors:** Xinyu Ye, Xiaodong Ma, Ziyi Pan, Zhe Zhang, Hua Guo, Kâmil Uğurbil, Xiaoping Wu

## Abstract

**Purpose:** to propose a two-step non-local principal component analysis (PCA) method and demonstrate its utility for denoising diffusion tensor MRI (DTI) with a few diffusion directions.

**Methods:** A two-step denoising pipeline was implemented to ensure accurate patch selection even with high noise levels and was coupled with data preprocessing for g-factor normalization and phase stabilization before data denoising with a non-local PCA algorithm. At the heart of our proposed pipeline was the use of a data-driven optimal shrinkage algorithm to manipulate the singular values in a way that would optimally estimate the noise-free signal. Our approach’s denoising performances were evaluated using simulation and in-vivo human data experiments. The results were compared to those obtained with existing local-PCA-based methods.

**Results:** In both simulation and human data experiments, our approach substantially enhanced image quality relative to the noisy counterpart, yielding improved performances for estimation of relevant DTI metrics. It also outperformed existing local-PCA-based methods in reducing noise while preserving anatomic details. It also led to improved whole-brain tractography relative to the noisy counterpart.

**Conclusion:** The proposed denoising method has the utility for improving image quality for DTI with reduced diffusion directions and is believed to benefit many applications especially those aiming to achieve quality parametric mapping using only a few image volumes.

## Introduction

Diffusion MRI (dMRI) provides a noninvasive way to detect tissue microstructure information by probing the Brownian motion of water molecules [1-2]. It is widely used in clinical diagnosis and neuroscience studies to measure the changes in diffusive properties of tissue micro-environment and connective features of the neural fibers [3-6]. However, one major challenge for dMRI is its intrinsically low signal-to-noise ratio (SNR) which hinders data acquisition with high resolution and/or high diffusion weighting [7-8]. It is shown that the low SNR resulting from pushing the resolution and/or diffusion weighting can reduce the accuracy of estimation of crossing fibers [9]. One way to maintain sufficient SNR is to acquire multiple scans for averaging. However, this will increase the scan time, which is undesirable for human studies. Thus, it is important to develop a postprocessing method (such as denoising) to improve SNR for dMRI without increasing the scan time.

Over the past decades, researchers have proposed various image denoising methods for filtering dMRI data before downstream diffusion analysis. This includes non-linear smoothing[10-12], non-local means[13], wavelet-domain filtering[14], principal component analysis (PCA) based approaches[15-16], among many others . A comprehensive review on denoising methods suitable for dMRI can be found in Ma et al[17]. Most notably, image denoising methods based on local PCA[15-20] (L-PCA) has been drawing increasing attention in recent years due in large to their demonstrated effectiveness in improving image quality for dMRI data.

L-PCA methods aim to denoise 4D dMRI series (3D space + 1D diffusion direction) by manipulating singular values obtained from the singular value decomposition (SVD) of local data patches. Local data patches for individual image voxels of interest can be obtained by sliding a kernel in space across the image field of view (FOV) and considering local data across all diffusion direction volumes. For each image voxel in space, this results in a local 4D data patch centered on the image voxel under consideration. The local 4D data patch is then used to form a Casorati matrix with which to estimate the underlying noise-free signal for every voxel within the 4D data patch. The estimation of underlying noise-free signal can be fulfilled using optimal singular value thresholding or shrinkage, assuming that the associated Casorati matrix is of low rank. As a result, multiple estimates of underlying noise-free signals are obtained for every image voxel, which can be aggregated (e.g., using weighted averaging) on a voxel-by-voxel basis when using a patch-based approach to form the final denoised 4D diffusion volumes.

The usefulness of L-PCA methods has been demonstrated for denoising both magnitude[15-19] and complex-valued[20-21] dMRI data. Although meant to remove Gaussian noise, L-PCA methods are shown effective for denoising magnitude data with Rician noise especially when SNR is relatively high[18]. However, when SNR becomes low (e.g., under ∼5), directly using L-PCA methods in the magnitude domain (i.e., to denoise magnitude data) would not work as well because the Rician distribution starting to substantially deviate from the Gaussian counterpart. In this case, it is shown[17] that converting the Rician data into Gaussian-like data via variance stabilizing transformation (VST)[22] can help improve the denoising performances. For data with further reduced SNR (e.g., when pushing the resolution to the limit), it is best practice to denoise in the complex domain (i.e., to denoise complex-valued data which are usually Gaussian distributed) to leverage the L-PCA methods and maximize their denoising performances. Previous studies]17,20] have shown that denoising in the complex domain outperformed that in the magnitude domain with or without VST.

We note that the efficacy of L-PCA methods heavily rely on sufficient data redundancy available in the diffusion dimension to ensure the low rankness of individual Casorati matrices when formed using local data patches. This is usually the case when acquiring dMRI with many image volumes or diffusion encoding directions such as in high angular resolution diffusion imaging (HARDI)[23]. However, when acquiring dMRI with few diffusion encoding directions such as in diffusion tensor imaging (DTI)[24], using L-PCA methods is likely to lead to sub-optimal denoising performances, due in large part to the lack of data redundancy in the diffusion direction dimension. In this case, a better choice would be to use a non-local PCA (NL-PCA) approach[25-27] where similar non-local data patches are identified and used to form the Casorati matrix to promote its low rankness by including the data similarity available in space.

A key procedure involved in NL-PCA approaches is patch selection or patch matching based on image self-similarity. In patch selection, non-local patches that have similar image content to the reference patch (i.e., the local 4D data patch of the image voxel under consideration) are identified in a predefined searching zone (e.g., across the entire image FOV). The similarities of non-local patches to the reference patch can be evaluated by calculating their Euclidian distances to the reference patch. The efficacy of Euclidian distances in determining similar non-local patches has been demonstrated for existing denoising methods[25-26] aiming to capitalize on grouping similar non-local patches. However, when denoising images with low SNR, the calculation of Euclidian distances may be inaccurate due in large to high noise levels, degrading the patch selection. In this case, it is shown[27] that patch selection can be effectively improved by basing the calculation of Euclidian distances on pre-denoised “guide” images.

Here, we propose an improved NL-PCA approach for denoising complex-valued 4D timeseries with a few image volumes and demonstrate its effectiveness in improving image quality for DTI. Preliminary results are reported in the format of conference abstracts[28-29]. Our improved NL-PCA approach is devised to have two steps, with step 1 aiming to provide the pre-denoised images that are used to guide the patch selection for step 2 to fulfill the task of NL-PCA denoising. We evaluated the efficacy of our NL-PCA approach using both simulation at 3 Tesla (3T) and in-vivo human data experiments at 7T. The human brain data were collected using a commercially available 32-channel receive RF head coil (Nova 1Tx/32Rx). In both simulation and in-vivo human data experiments, our results showed that our improved NL-PCA approach largely improved the image quality for DTI with nine image volumes when compared to the noisy counterpart, outperforming existing L-PCA methods.

## Method

We implemented the proposed non-local PCA denoising method in MATLAB (The Mathworks Inc., Natick, MA). The source code will be made publicly available at https://github.com/ye135246/Non_local_denoise. To demonstrate the efficacy of our proposed method, both simulation and in-vivo human data experiments were carried out where the proposed method was applied to DTI data. The results were compared to those obtained without denoising, as well as those using two L-PCA approaches: MPPCA[18] (applied to magnitude images as originally proposed) and NOise reduction with DIstribution Corrected (NORDIC) PCA[20]. For both simulation and in-vivo data experiments, MPPCA denoising was performed by using the implementation in the MRtrix3 package [30], whereas NORDIC was implemented based on the code shared at https://github.com/SteenMoeller/NORDIC_Raw.

### Two-step non-local PCA denoising

We performed image denoising in the complex domain (i.e., to denoise complex-valued images) by devising a two-step non-local PCA method (Fig. 1). Our proposed two-step method started with data preprocessing where the input noisy complex-valued images underwent phase stabilization (i.e., to remove background phases and direction-dependent phases) and g-factor normalization (i.e., to account for spatial variations due to g-factor). Phase stabilization was performed using the same way as in NORDIC. G-factor normalization was achieved by dividing the phase stabilized data by an estimated g-factor map. The g-factor map was estimated by applying the MPPCA approach [18] (with a sliding spatial kernel of size 5×5×5) only to the real part of the phase stabilized noisy data.

**Fig. 1.**
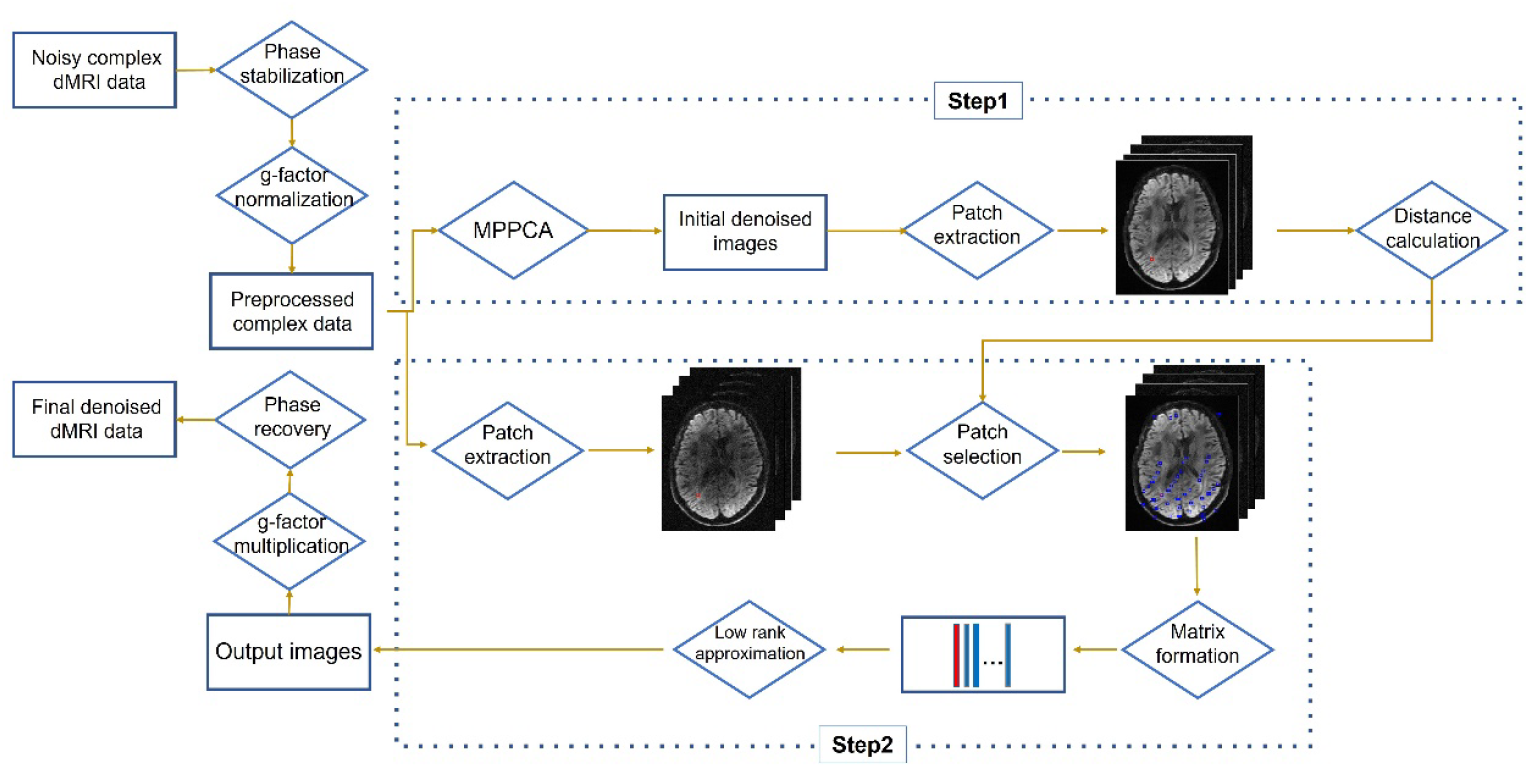
Flowchart of our proposed 2-step non-local PCA denoising method. Original noisy complex-valued diffusion MRI (dMRI) data are preprocessed with phase stabilization (i.e., to remove smooth phases) and g-factor normalization (i.e., to divide by g-factors) before being fed into a 2-step denoising pipeline. In step 1, Marchenko-Pastur PCA (MPPCA) is used to denoise noisy images for improved patch selection. In step 2, similar non-local patches are selected (per the distance calculation from step 1 to form a Casorati matrix with which to estimate low rank components using optimal singular value shrinkage (a PCA-based algorithm) for improved denoising. The red box refers to an example patch of interest and the blue boxes the selected similar patches. The output images of step 2 are further manipulated by multiplying with g-factors and restoring the smooth phases removed in phase stabilization, to produce the final denoised images.

The preprocessed complex-valued noisy images were then fed into a two-step denoising pipeline. In step 1, only the real part of the input images was considered and denoised with MPPCA, creating intermediate images with improved quality to facilitate patch selection. For each spatial location in the 3D space, a 4D patch (i.e., 3D space + 1D diffusion direction) was extracted and its similarity to every other non-local patch inside a prescribed neighborhood in space was evaluated by calculating the Euclidian distance. In step 2, the preprocessed complex-valued noisy images were denoised with a non-local PCA algorithm, for which patch selection was guided by the calculation of Euclidian distances conducted in step 1. Specifically, each noisy patch was grouped with Q similar non-local patches (i.e., the first Q non-local patches with the smallest Euclidian distances to the patch) to form the Casorati matrix. The value Q was adaptively updated. Here, we defined the similar patches as those whose distance was smaller than 1.25 × *d*_*min*_, where *d*_*min*_ is the smallest distance between all other patches with the selected patch. The Casorati matrix’s low-rank components were estimated using optimal singular value shrinkage [31] given its demonstrated denoising efficacy identified in our previous study[17]. Patch averaging similar to Ma et al. [17] was used to form the output images of step 2, which in turn were further postprocessed to form the final denoised complex-valued images by multiplying with g-factors and restoring the smooth phases that had been removed in phase stabilization. In both steps, 4D patches were formed by sliding a spatial kernel of size 5×5×5 across the image FOV with a step size of 2.

### Simulation experiment

We started with a simulation experiment to demonstrate the efficacy of our proposed denoising method for DTI. Synthetic data were generated as follows. Noise-free magnitude data were simulated based on part of a single subject’s 3T Human-Connectome-Project (HCP) preprocessed dMRI data [32]. Specifically, a total of 108 images (including 18 b=0 and 90 b=1000 s/mm^2^ images) obtained at 1.25-mm resolutions was used to fit a tensor model with fsl’s *dtifit* routine. The obtained tensor in turn was used in a DTI signal model [33] to synthesize noise-free magnitude data comprising a total of nine images (including one b=0 and eight b=1000 s/mm^2^ images). Then noise-free magnitude data were manipulated with second-order smooth phase variations to synthesize the noise-free complex-valued data.

Noise-corrupted DTI data with different noise levels were synthesized by adding 3D spatially-varying Gaussian noise to noise-free complex-valued data (serving as a gold standard). Specifically, spatially-varying Gaussian noise was added to both real and imaginary parts of noise-free data. The noise level was defined by the ratio of the maximum standard deviation (std) of the underlying Gaussian noise and the maximum signal intensity of the noise-free b0 image. Noisy data with noise levels ranging from 1% to 10% in steps of 1% were created.

For each noise level, the denoising performance of our proposed method was evaluated by comparing the final denoised images to the gold standard images in the magnitude domain. This was done by calculating two image quality assessment metrics: Peak SNR (PSNR) and Structure Similarity (SSIM) [34]. Both PSNR and SSIM values were calculated by considering all images including the b0 image.

Furthermore, diffusion analysis was conducted to investigate how our proposed denoising method would improve DTI. For this, diffusion tensor model was fit for the denoised magnitude images using FSL’s *dtifit* routine to obtain DTI metrics including fractional anisotropy (FA) and mean diffusivity (MD). For each DTI metric, the normalized root mean square error (nRMSE) was evaluated to measure the deviation from the gold standard which was obtained by fitting the diffusion tensor model to the noise-free images. In all cases, the results were compared to those obtained from the noisy counterparts, and those from the denoised images using MPPCA and NORDIC (both with a sliding kernel of size 5×5×5).

### In vivo data experiment

We also performed in-vivo human data experiment to evaluate the denoising performances of our proposed method. For this, brain dMRI data were collected with higher spatial resolutions on a 7T Siemens Terra scanner (Siemens, Erlangen, Germany) equipped with a body gradient (capable of 80 mT/m maximum gradient strength and 200 T/m/s maximum slew rate). One healthy adult who signed a consent form approved by the local Institutional Review Board was scanned using the commercial Nova 32-channel receive coil. Multiband-accelerated whole-brain DTI data were acquired at 0.9-mm isotropic resolutions using single-shell q-space sampling (b=1500 s/mm^2^) as in the 7T HCP protocol [35]. Other relevant imaging parameters were: 2-fold multiband acceleration, 3-fold in-plane acceleration, and TR/TE=7000/70 ms. The dataset comprised 20 averages in total, each with nine image volumes (corresponding to one b0 and eight diffusion directions). Each average consisted of two runs with opposed phase encodes: One with Anterior-Posterior (AP) and the other PA phase encodes to allow for correction of geometric distortions in the subsequent diffusion image preprocessing.

All original dMRI data were reconstructed offline in MATLAB using a custom reconstruction program. Multi-channel images of individual receive channels were first reconstructed using an improved 3D GRAPPA algorithm (involving a new 2-stage N/2 ghost correction and the GRE single-band reference for improved reconstruction) [36], and were then combined via adaptive coil combination [37] to form the final noisy complex-valued diffusion images.

We randomly chose a single average for denoising. The denoising was done at the run level, i.e., the two runs (of nine images each) were denoised separately as in Ma et al[17]. The background voxels were excluded in the denoising processes for improved computation efficiency.

The denoised single average images were then preprocessed in the magnitude domain following the HCP pipeline [38] to correct for head motion and EPI distortion. This was done by using FSL’s *topup* and *eddy* routines[39-40]. Further, the corrected images were registered to the volunteer’s native structural volume space defined by T1-weighted (T1w) and T2w images at 0.7-mm isotropic resolutions as in our previous studies[17,41-42].

Finally, we performed diffusion analysis using the denoised single average images to derive DTI metrics including FA and MD as in the simulation experiment. The results were compared to those derived from the dataset with 20 averages (serving as a reference). Note that all preprocessed images of the 20 averages were treated as independent volumes with no prior averaging in the image domain when used to fit a diffusion tensor model. DTI analysis results for the same single average without denoising, denoised using MPPCA and denoised using NORDIC were also obtained for comparison.

To investigate how our proposed denoising method would improve fiber tracking, we created whole-brain tractography using the DTI analysis results for the denoised single average images. This was done in the DSI studio (https://dsi-studio.labsolver.org/) using a deterministic fiber tracking algorithm [43] with augmented tracking strategies [44] for improved reproducibility. The anisotropy threshold was randomly selected. The angular threshold was randomly selected from 15 to 90 degrees. The step size was set to voxel size. Tracks with a length shorter than 30 or longer than 200 mm were discarded. A total of one million seeds were placed. The resultant whole-brain tractography was compared to that obtained using the same single average but without denoising.

## Results

### Simulation

Our proposed method substantially improved the image quality relative to the noisy counterparts across all noise levels under testing (Fig. 2), increasing both PSNR and SSIM values. Quantitatively, PSNR increased by 10.8% (∼53 vs ∼48) at 1%, by 15.9% (∼49 vs ∼42) at 2%, by ∼19.6% (∼46 vs ∼38) at 3%, by 22.4% (∼44 vs ∼36) at 4%, by ∼24.7% (∼42 vs ∼34) at 5%, and by 26.6% (∼41 vs ∼32) at 6% noise level, by 28.1% (∼40 vs ∼31) at 7%, by ∼29.3% (∼39 vs ∼30) at 8%, and by 30.4% (∼38 vs ∼29) at 9%, by 31.3% (∼37 vs ∼28) at 10% noise level whereas SSIM increased by 0.6% (∼0.99 vs ∼0.99) at 1%, by 2.6% (∼0.99 vs ∼0.97) at 2%, by ∼5.9% (∼0.99 vs ∼0.93) at 3%, by 10.2% (∼0.98 vs ∼0.89) at 4%, by ∼15.1% (∼0.97 vs ∼0.85) at 5%, and by 20.3% (∼0.96 vs ∼0.80) at 6%, by 25.7% (∼0.95 vs ∼0.76) at 7%, by ∼30.8% (∼0.94 vs ∼0.72) at 8%, and by 35.8% (∼0.92 vs ∼0.68) at 9%, by 40.3% (∼0.91 vs ∼0.65) at 10% noise level. Our proposed method also outperformed both MPPCA and NORDIC in improving image quality especially at higher noise levels, increasing PSNR by up to ∼16.7% (∼37 vs ∼31) relative to MPPCA and by up to ∼12.8% (∼39 vs ∼34) relative to NORDIC while increasing SSIM by up to ∼9.5% (∼0.91 vs ∼0.83) relative to MPPCA and by up to ∼9.9% (∼0.91 vs ∼0.83) relative to NORDIC.

**Fig. 2.**
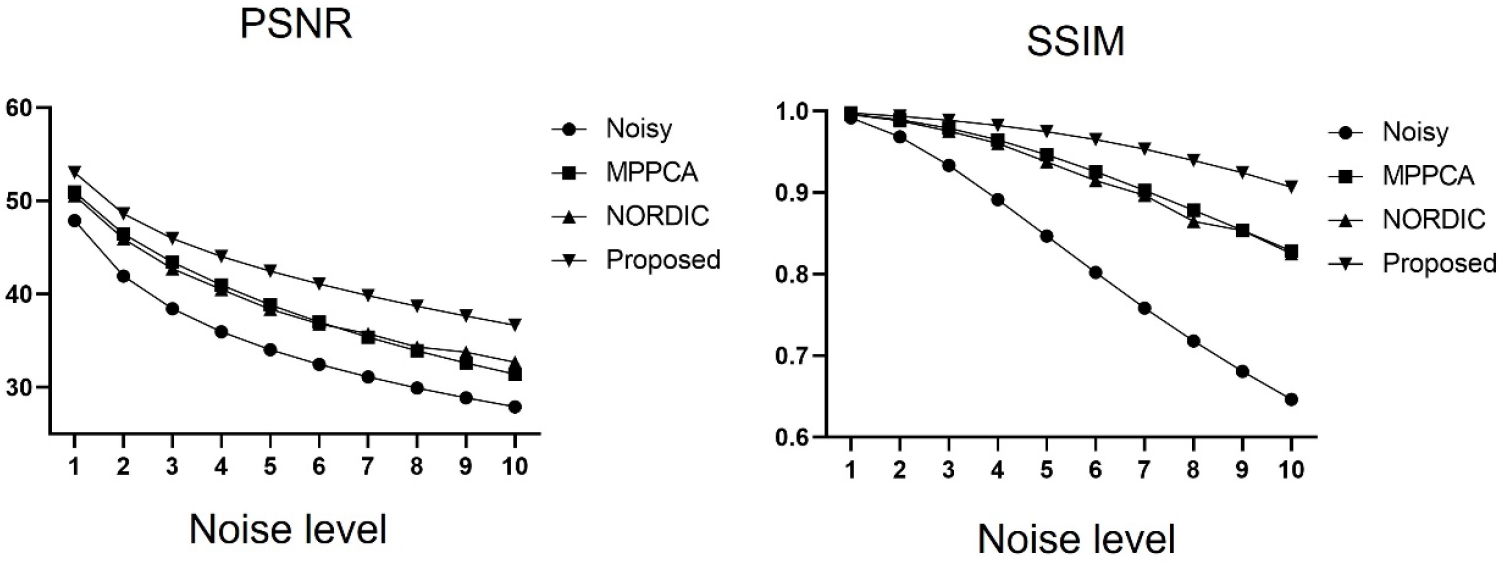
Simulation experiment: comparing denoising performances of our proposed 2-step non-local PCA (Proposed) vs existing local PCA approaches including MPPCA (applied in the magnitude domain as originally proposed) and NOise Reduction with DIstribution Corrected PCA (NORDIC) (g-factor normalization + phase stabilization + MPPCA), in terms of overall peak SNR (PSNR) in dB and structural similarity index measure (SSIM) as a function of the noise level (%). PSNR and SSIM values for noisy images are also shown for comparison. In all cases, both overall PSNR and SSIM values were calculated by considering b0 and diffusion weighted images. Synthetic noise-free data (including one b0 and 8 b=1000 s/mm^2^ images) serving as a gold standard were created based on one subject’s 3T dMRI data from the original young adult Human Connectome Project (HCP) and the noisy data were created by adding spatially varying Gaussian noise to the noise-free data. Note that the use of the proposed denoising method outperformed all the local PCA approaches under consideration, substantially improving the image quality (especially for higher noise levels).

The improvement in image quality with our proposed method was further confirmed by examining the diffusion weighted images at two representative noise levels of 4% and 6% (Fig. 3). Our proposed method substantially reduced the noise compared to the noisy counterpart, restoring fine brain structure across the brain. It also reduced the noise more effectively than both MPPCA and NORDIC, bringing the image quality closer to that of the gold standard especially at a higher noise level.

**Fig. 3.**
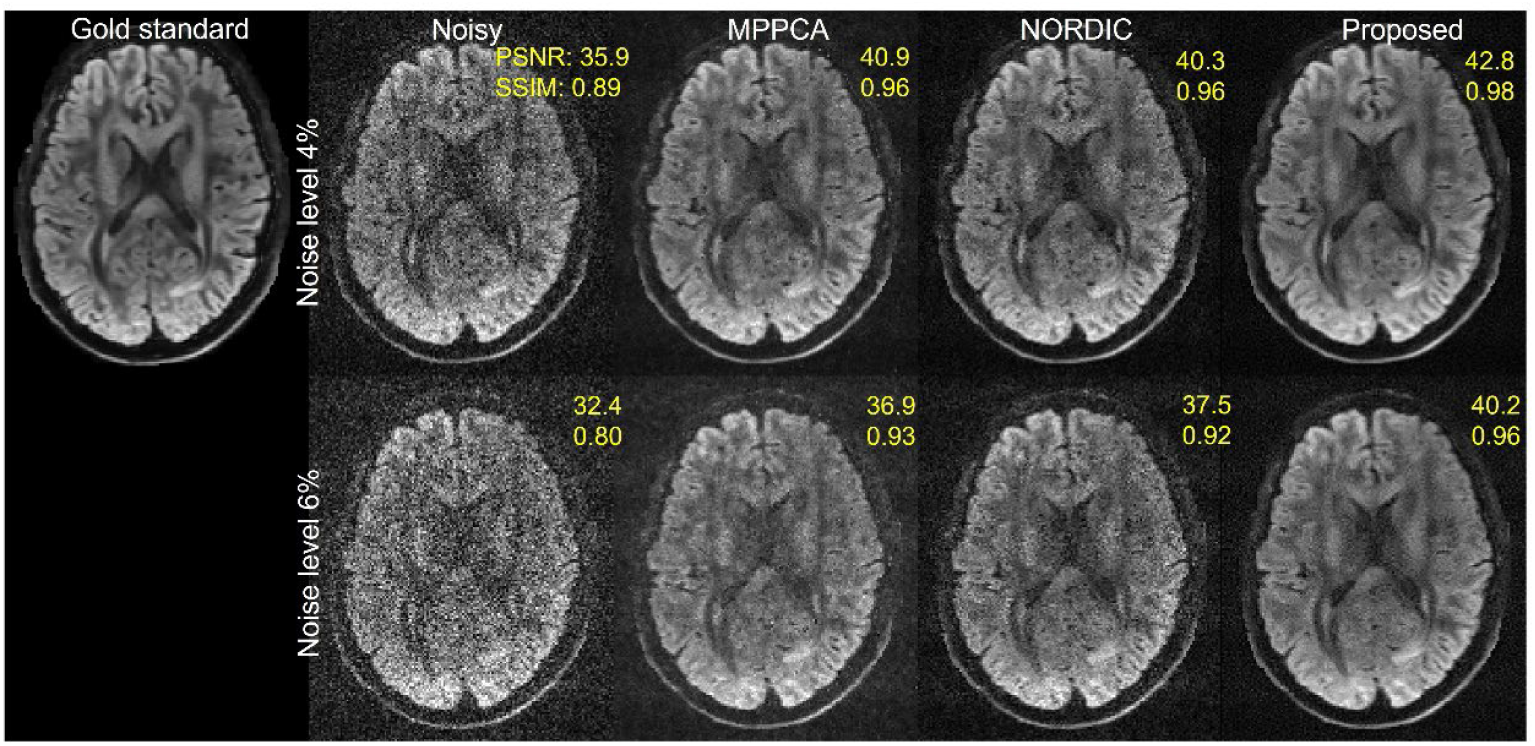
Simulation experiment: comparing denoised diffusion weighted images at two representative noise levels (4% and 6%) obtained using MPPCA, NORDIC, and our proposed method, alongside the corresponding gold standard and noisy images. In each case, shown are diffusion weighted images of one representative diffusion direction in the axial view. The numbers reported are respective PSNR (in dB) and SSIM values calculated relative to the gold standard. Note that the use of our proposed method outperformed both MPPCA and NORDIC in improving image quality, yielding highest PSNR and SSIM values at either noise level.

The improvement in image quality with our proposed method relative to the noisy images translated into increased performances for estimation of DTI metrics at all noise levels under consideration (Fig. 4), reducing nRMSE values for both FA and MD. Quantitatively, nRMSE decreased by 31.9% (∼5.7% vs ∼8.4%) at 1%, by 42.8% (∼8.9% vs ∼15.6%) at 2%, by ∼50.8% (∼11.4% vs ∼23.1%) at 3%, by 55.8% (∼13.2% vs ∼29.8%) at 4%, by ∼59.4% (∼14.4% vs ∼35.4%) at 5%, and by 61.5% (∼15.4% vs ∼40.0%) at 6%, by 62.4% (∼16.4% vs ∼8.69%) at 7%, by 62.2% (∼17.6% vs ∼46.5%) at 8%, by ∼61.3% (∼18.9% vs 48.8%) at 9%, by 59.5% (∼20.5% vs ∼50.6%) at 10% noise level for FA, whereas by 22.2% (∼3.0% vs ∼3.9%) at 1%, by 33.1% (∼5.1% vs ∼7.6%) at 2%, by 41.1% (∼6.7% vs ∼11.4%) at 3%, by 46.2% (∼8.1% vs ∼15.0%) at 4%, by ∼49.3% (∼9.3% vs ∼18.4%) at 5%, and by 50.4% (∼10.6% vs ∼21.5%) at 6%, by 50.3% (12.0% vs ∼24.2%) at 7%, by 49.8% (∼13.3% vs ∼26.6%) at 8%, by ∼48.8% (∼14.7% vs ∼28.8%) at 9%, and by 47.4% (∼16.2% vs ∼20.4%) at 10% noise level for MD. Our proposed method also outperformed both MPPCA and NORDIC in improving estimation performances, reducing nRMSE for FA by up to ∼33.9% (∼17.6% vs ∼26.6%) relative to MPPCA at 8% noise level and by up to ∼33.6% (∼17.6% vs ∼26.5%) relative to NORDIC at 8% noise level while reducing nRMSE for MD by up to ∼36.5% (∼10.7% vs ∼16.8%) relative to MPPCA at 6% noise level and by up to ∼36.3% (∼10.7% vs ∼16.7%) relative to NORDIC at 8% noise level.

**Fig. 4.**
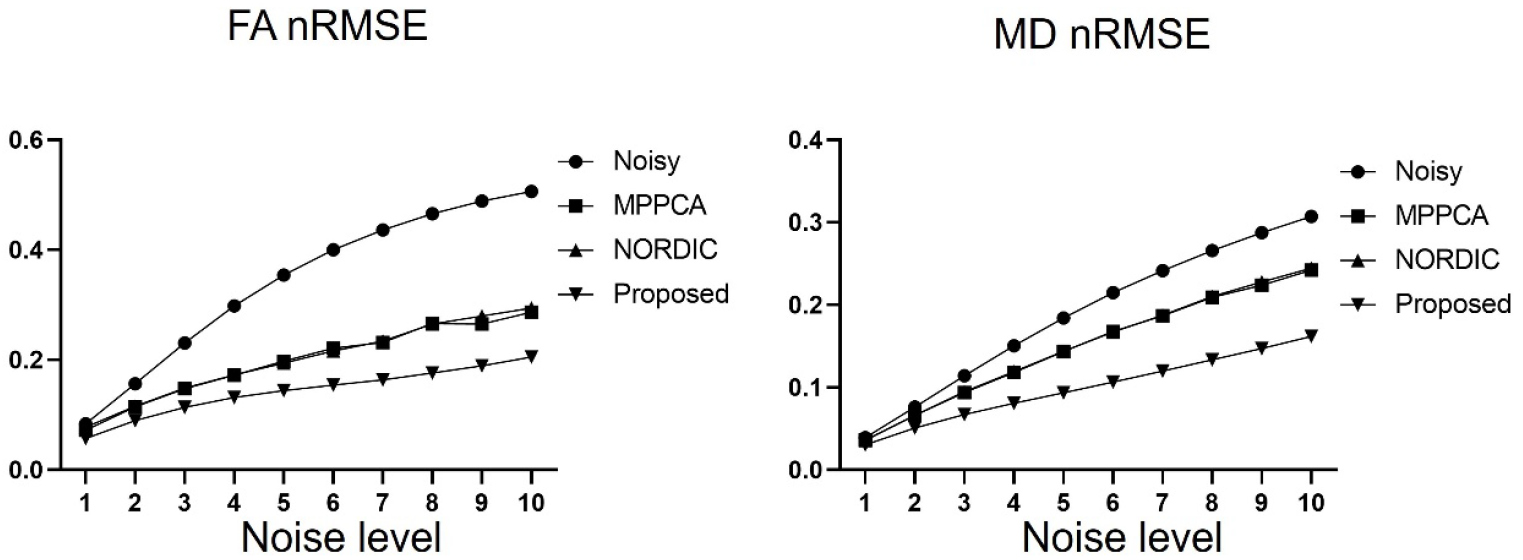
Simulation experiment: comparing denoising performances of our proposed 2-step non-local PCA (Proposed) vs existing local PCA approaches including MPPCA and NORDIC, in terms of normalized root mean squared error (nRMSE) for fractional anisotropy (FA) and mean diffusivity (MD) as a function of the noise level. The nRMSE values for noisy data are also shown for comparison. In all cases, FA and MD values were estimated by fitting a diffusion tensor model to the same data as for Fig. 2. All nRMSE values were calculated relative to what was obtained with the noise-free images serving as a gold standard. Note that the use of the proposed denoising method outperformed all the local PCA approaches under consideration, improving the performances for estimation of diffusion tensor metrics (especially for MD at higher noise levels).

The improvement in the estimation of DTI metrics using our proposed method, relative to the noisy images, was further confirmed by comparing FA and MD maps at two representative noise levels of 4% and 6% (Fig. 5). Our proposed method significantly enhanced both FA and MD maps, enabling visualization of fine brain structures. It also reduced the noise more effectively than MPPCA or NORDIC, bringing both FA and MD maps closer to those of the gold standard with less noise presented especially around the center of the brain.

**Fig. 5.**
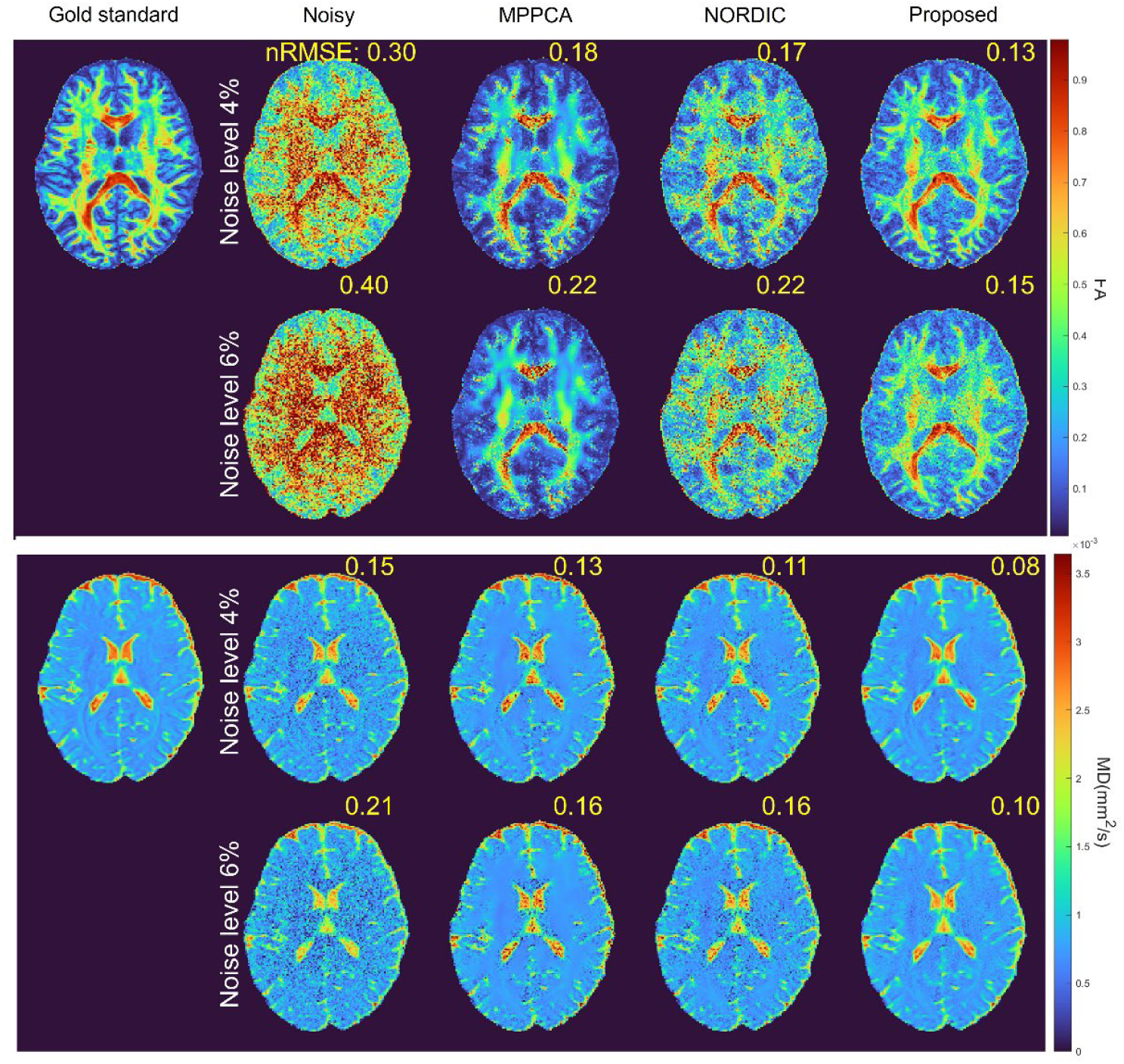
Simulation experiment: comparing estimation performances for fractional anisotropy (top panel) and mean diffusivity (bottom panel) maps at two representative noise levels (4% and 6%), obtained using MPPCA, NORDIC, and our proposed method, alongside what was obtained with corresponding gold standard and noisy images. Diffusion tensor metrics were obtained by fitting a diffusion tensor model to the same data as for Fig. 3. The numbers reported are respective normalized RMSE (nRMSE) values calculated relative to the gold standard. Note how the use of the proposed method outperformed both MPPCA and NORDIC, improving the estimation of fractional anisotropy and mean diffusivity maps with reduced noise at either noise level.

### In vivo experiments

Likewise, our proposed method largely improved the image quality for the in vivo dMRI at 0.9-mm isotropic resolutions (Fig. 6), enabling fine brain structures to be visualized across the whole brain when compared to the noisy counterpart. Visually, our method outperformed both MPPCA and NORDIC, improving noise reduction in general. Similar results to the simulation experiments were observed when comparing the DTI metrics (Fig. 7). Our proposed method outperformed both MPPCA and NORDIC, leading to FA values visually closer to the 20 averages while providing less underestimated MD values, especially in CSF as indicated by arrows. The improvement in performances for estimation of DTI analysis with our proposed method relative to the noisy counterpart was found to translate into an improvement in whole-brain fiber tracking (Fig. 8). The use of our proposed method effectively removed most of the spurious fibers observed for the noisy data, visualizing more clearly major fiber tracts (such as corpus callosum) and other short-range fibers across the entire brain.

**Fig. 6.**
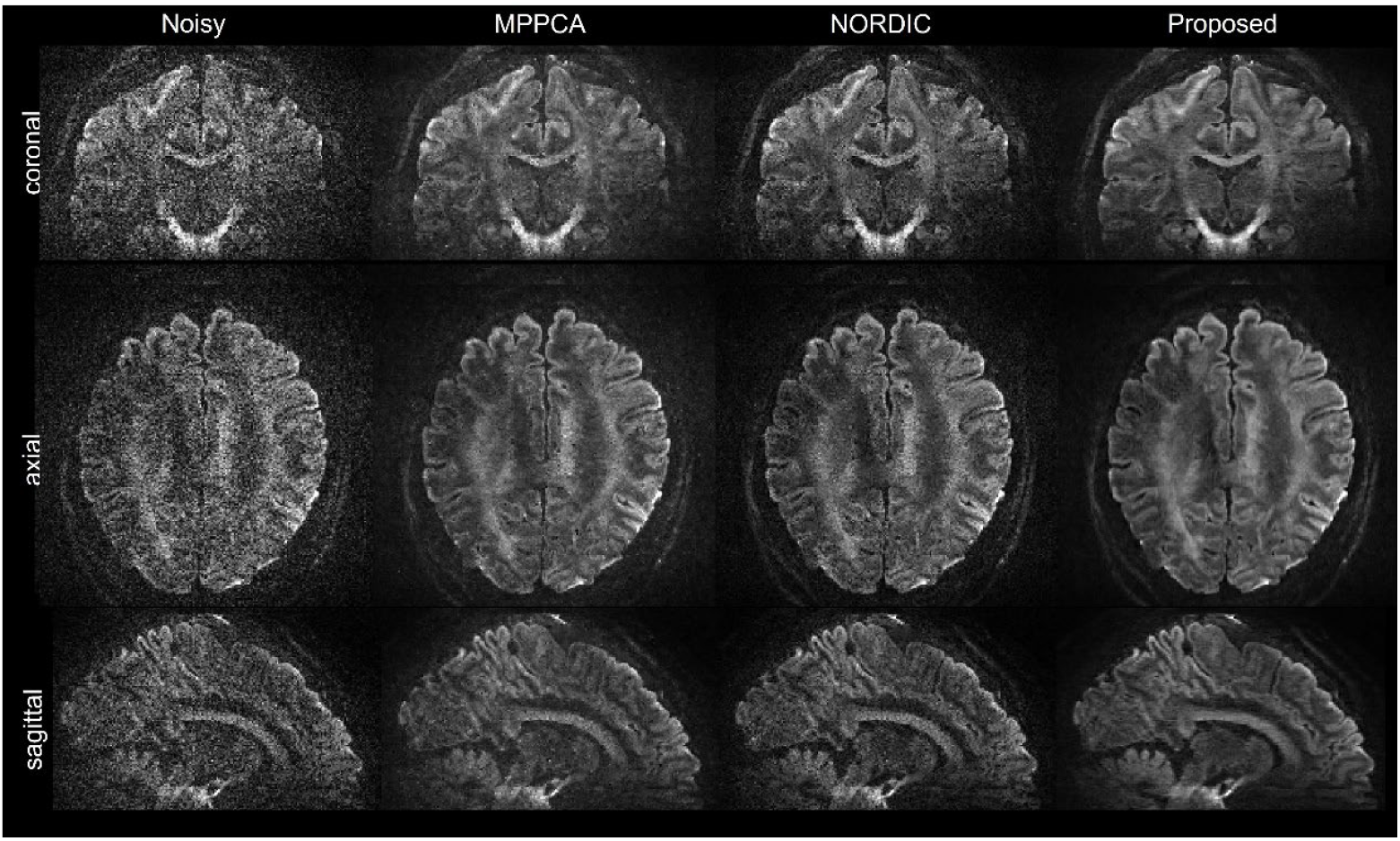
Human data experiment: comparing denoised diffusion weighted images at 0.9-mm isotropic resolution obtained using MPPCA (applied in the magnitude domain), NORDIC, and our proposed method, alongside the original noisy images for reference. In each case, shown are diffusion weighted images of one representative diffusion direction in the three orthogonal views. The original noisy images (including one b0 and 8 b=1500 s/mm^2^ images) were acquired with 2-fold slice acceleration, 3-fold in-plane acceleration, and TR/TE=7000/70 ms. Note how the use of our proposed method led to best image quality with least residual noise.

**Fig. 7.**
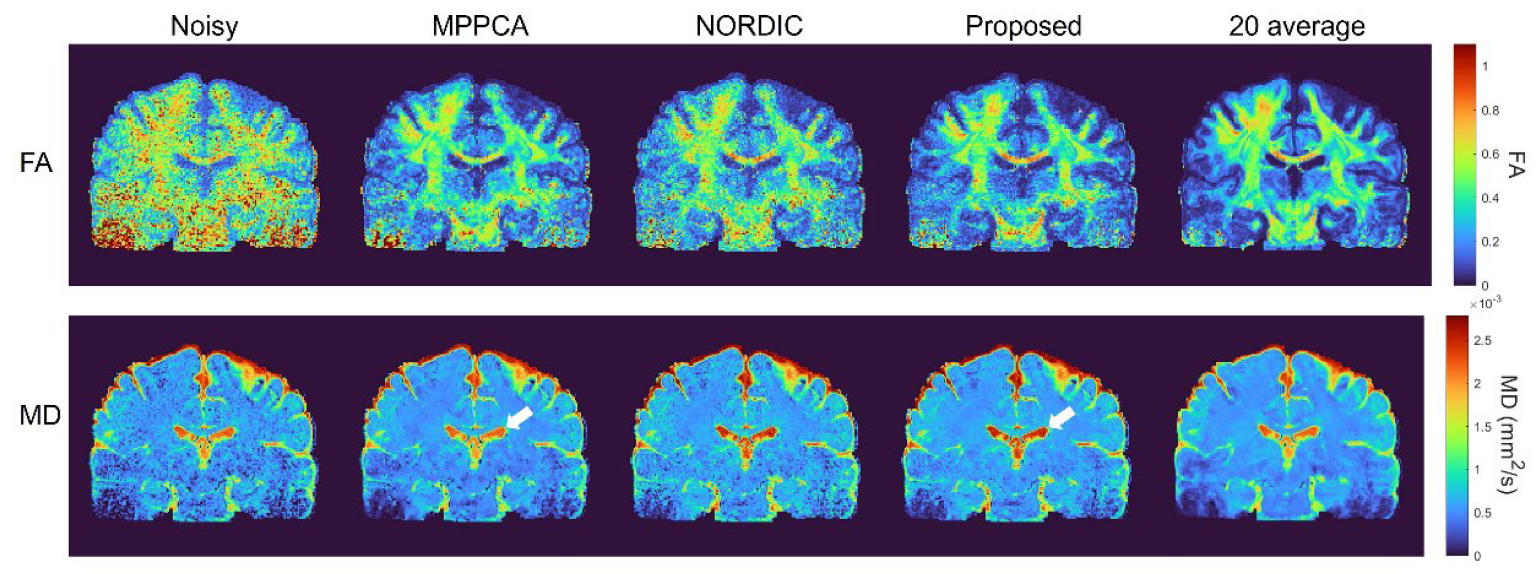
In-vivo human data experiment: comparing estimation performances for fractional anisotropy (FA) (top) and mean diffusivity (MD) (bottom), obtained using MPPCA, NORDIC, and our proposed method, alongside what was obtained with corresponding single average noisy images. In each case, diffusion tensor metrics were obtained by fitting a diffusion tensor model to the same single average data as for Fig. 6. The results for noisy data with 20 averages are also shown for reference. Note that the use of the proposed method outperformed both MPPCA and NORDIC, leading to FA values visually closer to 20 averages while providing less underestimated MD values, e.g., in CSF as indicated by arrows.

**Fig. 8.**
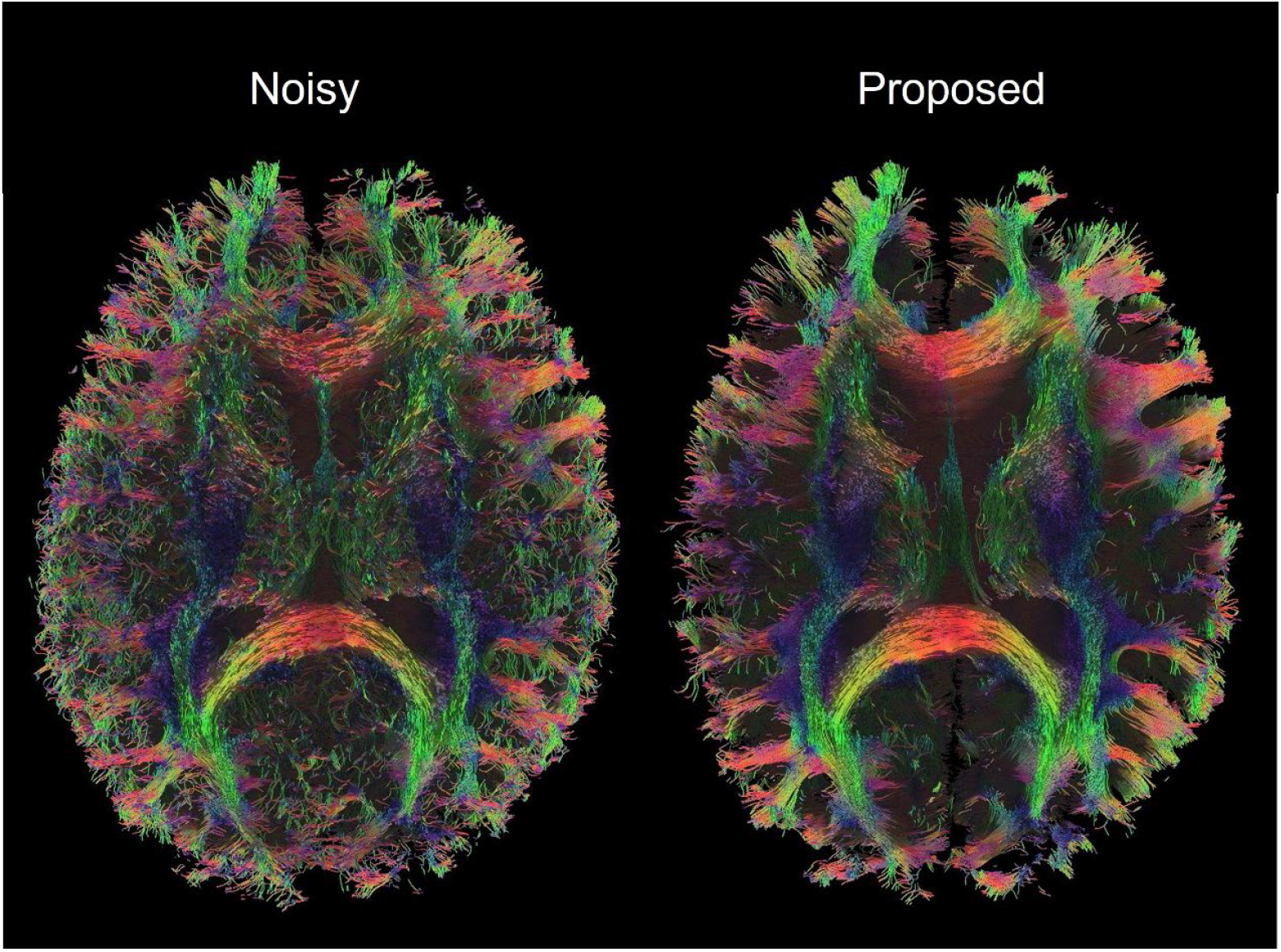
In-vivo human data experiment: demonstrating the utility of our proposed method for improving whole brain tractography in comparison to noisy images. In each case, shown is the color-coded tractography in the axial view. The tractography was obtained using a deterministic fiber tracking algorithm based on the same single average diffusion tensor analysis as for Fig. 7. Note how the use of the proposed method substantially improved the quality of whole brain tractography.

## Discussion

We proposed an improved non-local PCA approach suitable for denoising complex-valued dMRI images with a few diffusion directions. The efficacy of our proposed denoising approach was demonstrated for improving image quality of DTI using 3T synthetic and 7T in-vivo human data experiments. Critical to the effectiveness of our proposed approach was the implementation of a two-step denoising pipeline to ensure accurate patch selection even with high noise levels. Our proposed two-step denoising pipeline is coupled with data preprocessing in which original noisy data undergo g-factor normalization and phase stabilization before being denoised with a non-local PCA algorithm. At the heart of our proposed denoising pipeline is the use of a data-driven optimal shrinkage algorithm to manipulate the singular values in a way that would optimally estimate the noise-free signal. We validated our proposed denoising pipeline by conducting a simulation study where synthetic DTI data with nine image volumes were created based on a single subject’s data randomly chosen from the 3T HCP diffusion database. Our simulation results (Figs. 2-5) show that our proposed two-step non-local PCA approach can substantially improve image quality while enhancing the estimation of DTI metrics relative to the original noisy images especially at higher noise levels and that it can provide better denoising performances than existing local PCA methods. We also showcased the usefulness of our proposed denoising pipeline by collecting 7T whole-brain DTI at 0.9 mm isotropic resolutions with nine image volumes. Likewise, our in-vivo results (Figs. 6-8) show that our proposed approach can largely enhance image quality and down-stream diffusion analyses when compared to the original noisy counterpart, outperforming existing local PCA methods.

Our proposed denoising method is devised to start with data preprocessing in which the input noisy complex-valued dMRI data are preprocessed with phase stabilization and g-factor normalization. Phase stabilization is meant to eliminate volume to volume phase variations, thereby improving the low rankness in the subsequently created Casorati matrices. G-factor normalization is meant to remove the effect of spatially varying noises (due in large to the use of parallel imaging) so that the subsequently created Casorati matrices (based on the normalized data) are compatible with the employed optimal singular value shrinkage, a nonlinear singular value manipulation function derived assuming a natural data model[17,31] with zero-mean and unity-variance Gaussian noise. As in NORDIC[20], the image preprocessing is found to play a pivotal role in ensuring the denoising performances of our proposed two-step nonlocal PCA method. Indeed, our results (Fig. S1) from a pilot study show that including the image preprocessing would largely improve the denoising performances of our two-step denoising pipeline, increasing PSNR by up to ∼8% (36.6 vs 33.8 for when excluding the image preprocessing at a noise level of 7%) and eliminating the artifacts that would otherwise be observed especially at higher noise levels.

In the data preprocessing of our proposed denoising method, g-factor normalization is performed after phase stabilization. This is to capitalize on the improved low rankness of the subsequent Casorati matrices when formed using the phase-stabilized data. Moreover, the noise estimation necessary for g-factor normalization is implemented by applying MPPCA only to the real part of the phase-stabilized data to avoid the potential contamination from the imaginary part (which is considered to have little signal in it). Indeed, our results (Fig. S2) from a pilot study aiming to optimize noise estimation with MPPCA given the phase-stabilized data show that applying MPPCA to the real part would result in much more accurate noise estimation than applying MPPCA to the complex-valued data across the noise levels and kernel sizes under consideration. Our results (Fig. S2) also show that using a sliding kernel of size 5×5×5 (as chosen in the current study) would lead to best noise estimation performances especially at higher noise levels (≥ ∼4%) when compared to use of smaller or larger kernel sizes.

In the current implementation of our proposed two-step denoising method, we used MPPCA for two tasks: 1) to estimate noise in data preprocessing for g-factor normalization and 2) to denoise the preprocessed data in step 1 for creating the initial denoised images. However, these two tasks can also be fulfilled using alternative methods. For example, noise estimation in data preprocessing may be accomplished using a noise estimator tailored for Gaussian data, such as the one described in Gavish and Donoho[46] that works based on the median singular value of noise-corrupted data and the median of the Marcenko-Pastur distribution. The initial denoised images in step 1 may be obtained using other denoising approaches such as adaptive non-local means filtering[13]. Part of our future work is to investigate how the integration of alternative noise estimation and denoising methods in place of MPPCA would affect the denoising performances of our two-step denoising pipeline.

The results from our simulation experiments (Fig. 2) suggest that the use of our proposed two-step denoising method can effectively remove the negative impact of thermal noise, improving the image quality of a single average noisy data. To find out how many averages this improvement in image quality would be comparable with, we performed additional simulation study where the image quality of a single average denoised using our proposed method was compared to those obtained by averaging. Our results (Fig. S3) comparing PSNR and SSIM at various noise levels show that the use of our proposed method to denoise a single average would improve image quality to a level comparable to what is attainable with 10-15 averages especially at noise levels greater than ∼2%.

In this study, we demonstrated the utility of our proposed two-step non-local PCA denoising method for improving DTI image quality by considering data acquisition with nine image volumes. When denoising in the complex domain, the results (Figs. 2 and 3) from our simulation experiment show that our proposed method can outperform a local PCA alternative such as NORDIC, increasing both PSNR and SSIM values over a wide range of noise levels. This improvement in denoising performances is believed to stem from the ability of our proposed method to enhance the low rankness of the Casorati matrices by aggregating similar non-local data patches. However, the advantage of our proposed method over a local PCA approach is expected to reduce when denoising data acquisition with increased image volumes due in large to the increase in information redundancy in the time/diffusion direction becoming a dominant contributor to the low rankness of the Casorati matrices. To investigate how denoising performances would change with increasing image volumes when denoising in the complex domain, we performed additional simulation experiments where noisy complex data created with different image volumes were denoised using our proposed non-local PCA method and the results were compared to those obtained using NORDIC. Our data (Fig. S4) show that although the denoising performances of both methods appeared to increase with increasing image volumes, the improvement in denoising performances with our proposed method relative to NORDIC was found to decrease as image volumes increased, the increase in PSNR reducing from ∼10% (38.4 vs. 34.7 for 9 volumes) to ∼5% (42 vs. 39 for 27 volumes) and the increase in SSIM reducing from ∼5% (0.97 vs. 0.92 for 9 volumes) to ∼1% (0.98 vs. 0.97 for 27 volumes) when going from 9 to 27 image volumes.

One limitation of this study is that our proposed two-step non-local PCA denoising method in its current implementation has limited computation efficiency. This can lead to a long computation time due in large to the distance calculation needed for patch selection in step 1 and singular value decomposition of large-scale Casorati matrices needed for low rank approximation in step 2. Part of our future work is to investigate how the computation efficiency may be improved using a more powerful computer or a GPU or both.

Although tested and evaluated here using data acquisition in healthy volunteers, our proposed non-local PCA denoising method should have a utility for clinical applications. For example, in a recent study, we managed to demonstrate how our proposed non-local PCA method can be used to enable a 1-minute high resolution whole brain diffusion MRI in a clinical setup at 7T[47]. Part of our future work is to study how our proposed non-local PCA denoising method would help improve diagnosis of neurological diseases by considering more patients and various pathologies.

## Conclusion

We have proposed an improved 2-step non-local PCA approach for denoising complex-valued diffusion data and demonstrated its utility for enhancing image quality of DTI with a few diffusion directions. The efficacy of our proposed denoising approach was illustrated using both simulation and in-vivo human-data experiments. Our results show that our proposed approach can largely improve image quality and estimation performances for DTI metrics (including FA and MD), when compared to the original noisy counterpart. Our results also show that our approach can reduce noise more effectively than existing local PCA methods, thanks to its ability to promote low rankness by integrating non-local similar patches. We believe that our proposed method will benefit many applications especially those aiming to achieve quality parametric mapping using only a few image volumes.

## Acknowledgement

The authors would like to thank Steen Moeller for stimulating discussion on PCA based denoising and his assistance with NORDIC during the early stage of this project. XW and KU and all work carried out at the University of Minnesota were supported in part by the National Institutes of Health of the United States under grants R01 NS136490, P41 EB027061, and U01 EB025144. The content is solely the responsibility of the authors and does not necessarily represent the official views of the National Institutes of Health.

## Supporting Figures

**Fig. S1.**
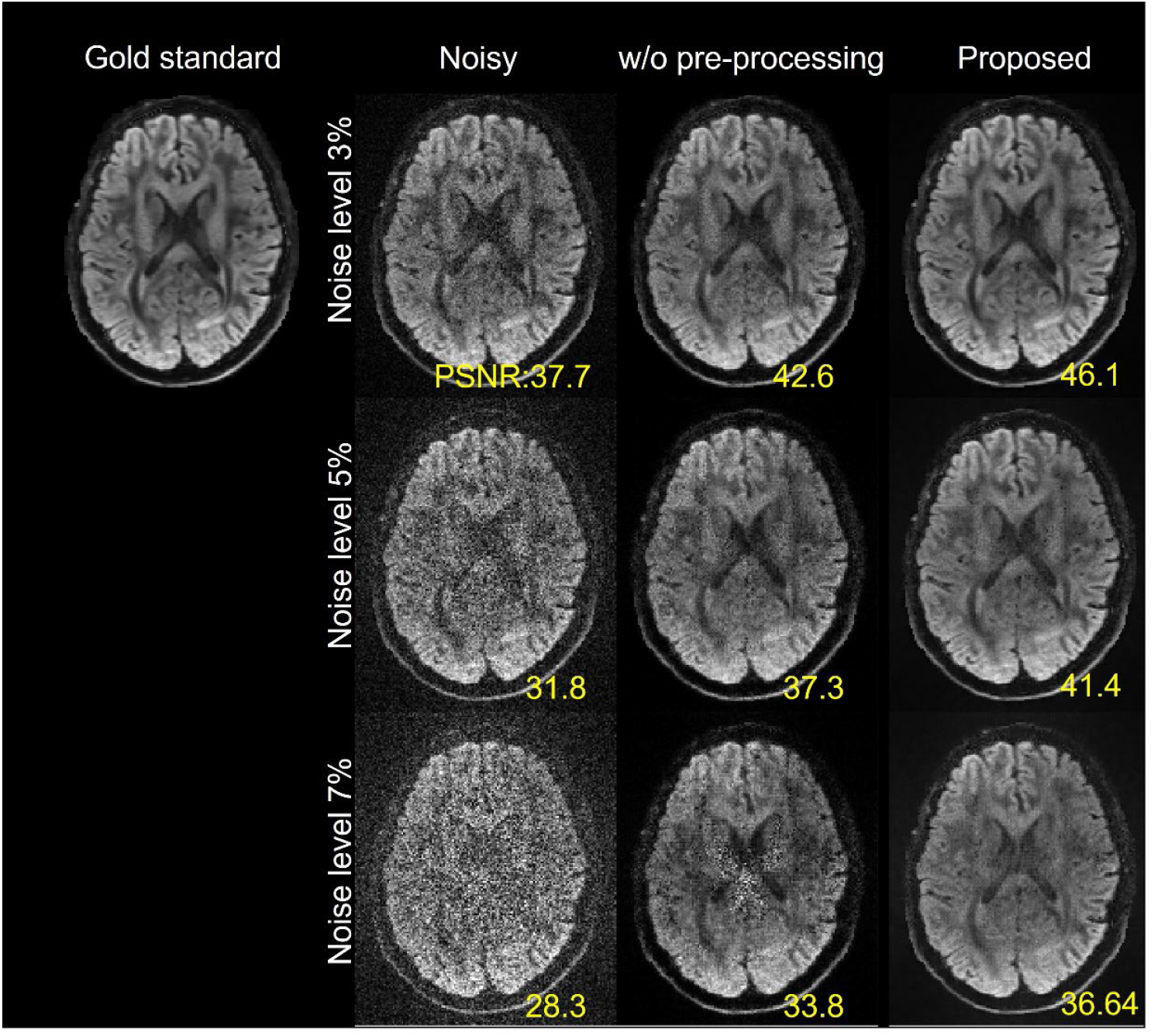
Simulation experiment: demonstrating the importance of data preprocessing (i.e., phase stabilization and g-factor normalization) in our proposed two-step non-local PCA denoising method. Shown are denoised images in a representative axial slice at three noise levels obtained using our proposed denoising method (Proposed) vs. those obtained using the same denoising pipeline but without data preprocessing (w/o pre-processing) vs. the original noisy counterparts, all in reference to the same gold standard as in Fig. 2. For quantitative analysis, PSNR values (in dB) are also reported. Note how including data preprocessing substantially improved the denoising performances especially at higher noise levels.

**Fig. S2.**
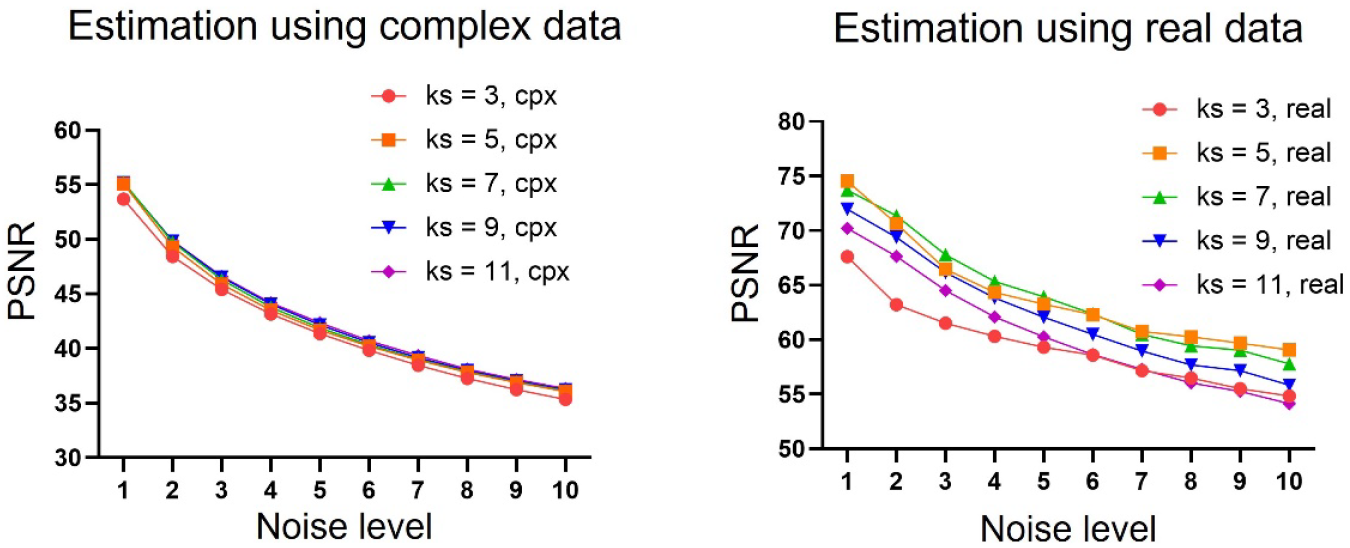
Simulation experiment: optimizing the noise estimation with MPPCA for g-factor normalization based on phase-stabilized complex-valued data. Shown are noise estimation performances (measured by PSNR) as a function of noise levels when applying MPPCA to the complex-valued data (left) vs. when applying MPPCA only to the real part of the complex-valued data (right). In both cases, noise estimation performances as a function of MPPCA kernel size (ks) are also shown at each noise level. PSNR values (in dB) at each noise level were calculated relative to the respective ground truth g-factor/noise map. The noise free dMRI data used to create the original noisy complex-valued data were the same as in Fig. 2. Note how applying MPPCA only to the real part of the phase-stabilized complex data with a 5x5x5 sliding kernel size led to best noise estimation performances especially at noise levels higher than ∼4%.

**Fig. S3.**
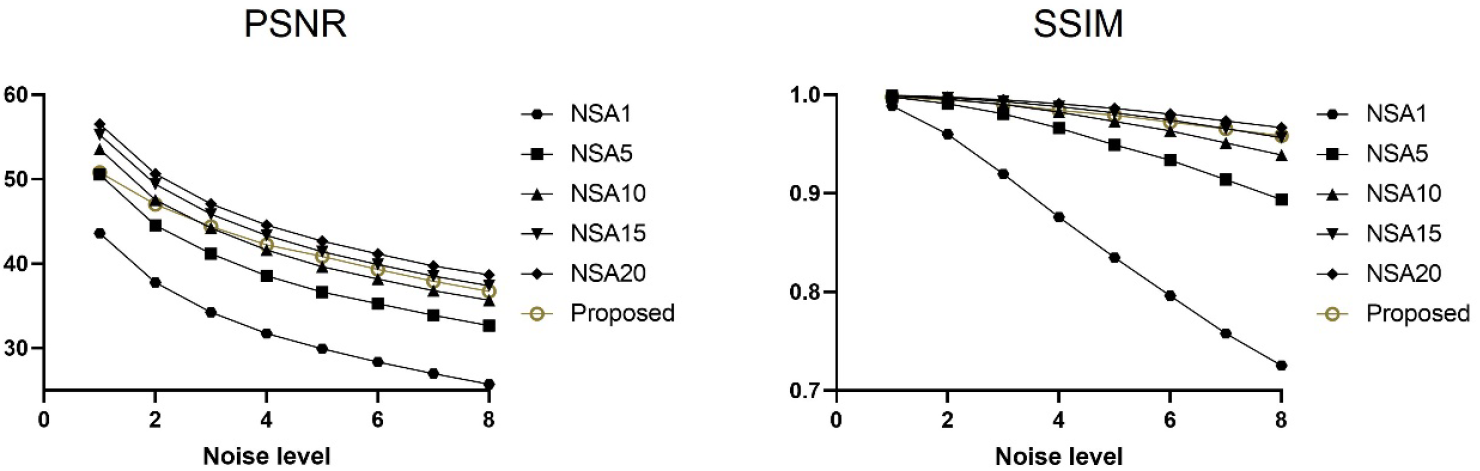
Simulation experiment: comparing our proposed denoising method with data averaging. Shown are PSNR (left) and SSIM (right) values as a function of noise levels for a denoised single average obtained with our proposed method (Proposed) vs. data averaging in the complex domain with different numbers of signal averages (NSA) and without denoising. Noisy complex data with 20 averages (NSA20) were created by drawing spatially varying Gaussian noise 20 times (from the same distribution) and adding them to the same noise-free complex data to mimic 20 scans. The noise-free complex-valued data used were the same as in Fig. 2. Note that denoising with our proposed non-local PCA method appeared comparable to data averaging with 10-15 averages, especially at noise levels greater than ∼2%.

**Fig. S4.**
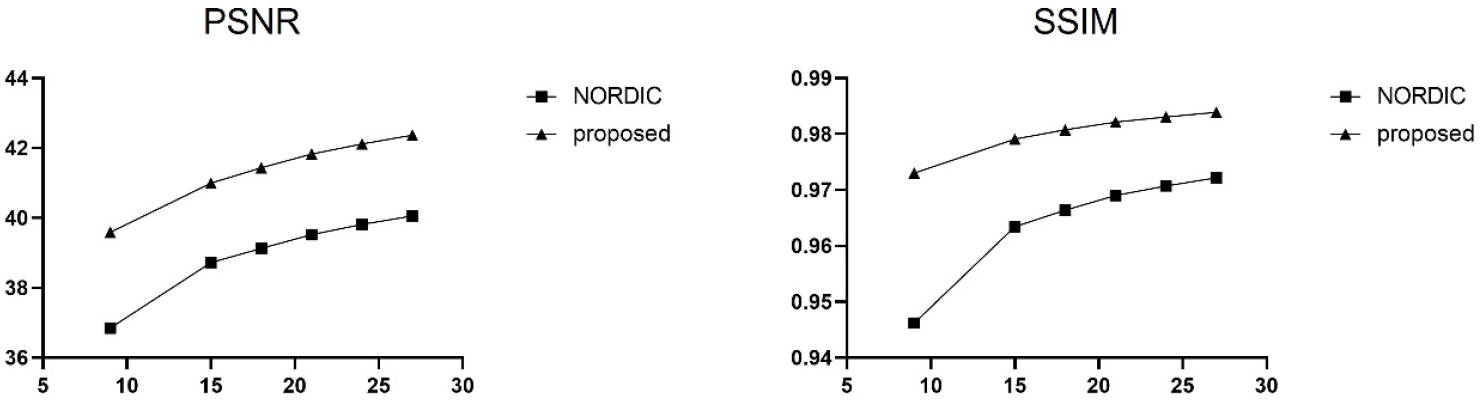
Simulation experiment: investigating how denoising performances would change with increasing image volumes/diffusion directions when denoising in the complex domain. Shown are PSNR (left) and SSIM (right) values as a function of the number, N, of image volumes (N ranging from 6 to 27 in steps of 3) when denoising with our proposed method (Proposed) vs with NORDIC at the noise level of 5%. Given N, synthetic noise-free data (including one b0 and N-1 b=1000 s/mm^2^ images) were created based on one subject’s 3T dMRI data from the original young adult Human Connectome Project (HCP) and the noisy data were created by adding spatially varying Gaussian noise to the noise-free data. Although the denoising performances of both our proposed method and NORDIC appeared to increase with increasing image volumes, note how the improvement in denoising performances with our proposed method relative to NORDIC decreased as image volumes increased especially when comparing SSIM values.

